# Time-Series Foundation Models for Cognitive Workload Classification using Eye-Tracking Data

**DOI:** 10.64898/2026.09.13.751282

**Authors:** Jordan Haag, Jose Gabriel Gonzalez Nunez, Brian J. Kirkwood, Gary L. Legault, Laura Brattain

## Abstract

Cognitive workload (CWL) assessment is relevant to a range of applications, such as monitoring driver fatigue and pilot attention, surgeon workload during complex procedures, and astronaut cognitive fatigue during long-duration missions. Eye-tracking datasets are generally small, which hinders the generalizability of the AI models. Time-series foundation models (TSFMs) have shown promise in mitigating this limitation as they are pretrained on large corpora and can be effectively fine-tuned with limited task-specific data. Although cognitive workload measurement has been explored in controlled settings, the ability of TSFMs to generalize to unseen individuals for CWL classification from eye-tracking data has not been studied. In this paper, two TSFMs, MOMENT and Moirai, were evaluated for CWL classification and compared against CNN, FFN, and LSTM baselines on two publicly available eye-tracking datasets. We used subject-level five-fold cross-validation in which data from each test subject were held out during training. We report accuracy, AUC, and F1-score with 95% confidence intervals. On the class-balanced EM-COGLOAD dataset, the pretrained TSFMs generalized markedly better to unseen subjects, with Moirai reaching 0.918 AUC and MOMENT 0.882 AUC, while the task-specific baselines remained near 0.70 AUC. On the class-imbalanced COLET dataset even though the performance of all the models decreased, MOMENT was the most robust (0.670 AUC). Both TSFMs were also the most stable across held-out subjects, indicating that pretrained representations generalize more consistently across individuals than task-specific models in data-scarce eye-tracking settings.

## I. Introduction

Cognitive workload (CWL) assessment is critical across multiple high-stakes disciplines. Elevated CWL over extended durations can lead to fatigue, stress, or distraction, which is directly relevant to driver safety [1], [2], the operational effectiveness of soldiers and pilots [3], and the performance of surgeons and astronauts [4]. CWL is considered a subjective experience relative to a specific task and therefore varies between individuals in similar situations [5]. One effective approach to CWL assessment is eye-tracking technology, which has demonstrated significant potential when paired with machine learning models [5]–[8]. Eye-tracking utilizes cameras to capture features such as saccades, gaze position, blink rate, and pupil diameter [9]. Considerable pre-processing is typically required to remove noise, separate fixations from saccades, and fill gaps caused by blinks [6], [7]. Ground-truth CWL labels are generally obtained directly from the experimental condition or through the NASA Task Load Index (NASA-TLX), which is a widely used, subjective, multidimensional assessment tool that rates perceived workload in order to assess a task [10].

The large amount of high-quality eye-tracking data needed to train ML models is often difficult to obtain. Custom classifiers are usually trained for a specific task [6]–[8] and do not generalize well to unseen data. Pre-trained on large and varied corpora [11], time-series foundation models (TSFMs) have shown promise in mitigating the limitations of small datasets. We hypothesize that TSFMs can learn effectively from small eye-tracking data without extensive data pre-processing. Commonly-used TSFMs include TimesFM [12], Chronos [13], and MOMENT [14]. More recently, Moirai [15] introduced an any-variate attention mechanism supporting variable context lengths and patch sizes.

In this paper, we evaluated two TSFMs, MOMENT-small and Moirai-small, on CWL classification from the EM-COGLOAD [7] and COLET [6] datasets. Our contributions are: (1) to our knowledge, the first evaluation of pretrained TSFMs for eye-tracking-based CWL classification using subject-level k-fold cross-validation, benchmarked against convolutional neural network (CNN), feedforward neural network (FFN), and long short-term memory (LSTM) baselines under a common statistical protocol; and (2) a comparison of how well pretrained TSFMs and task-specific models each generalize to unseen individuals, including their stability across held-out subjects. Each dataset is evaluated independently, as their channels represent different ocular measurements.

## II. Methods

### A. Data Description and Pre-processing

The first dataset, EM-COGLOAD, comprised eye-movement data from 75 participants who performed a visual-tracking task. It had binary CWL labels (high/low) based on the presence of a secondary task, a sampling rate of 160 Hz, and provided data from two pupils over four channels (left and right pupil vertical and horizontal positions). All four channels were used in this paper. We trimmed the samples to remove the first 14 seconds (calibration) and split them into 512-length training samples, excluding samples with missing entries [7]. The second dataset, COLET, consisted of gaze and pupil data from 47 participants across visual-search tasks of varying durations, sampled at 240 Hz and provided with continuous NASA-TLX workload scores [10]. We converted these scores into binary labels by thresholding at the midpoint of the scale, assigning scores of 0–49 to the low class and 50–100 to the high class. We used two channels of horizontal and vertical gaze position and two of left and right pupil diameter, and interpolated the data during blinks in the pupil-diameter channels using cubic splines for gaps below 500 ms [16]. We downsampled both datasets by half to make longer events fit within the 512-sample window length of the TSFMs. Table I summarizes dataset properties after downsampling. TSFMs process fixed-length tokens irrespective of sampling frequency (512 tokens correspond to either 6.4 s at 80 Hz or 4.3 s at 120 Hz), and hence we use and evaluate each dataset independently, not compared against the other [7]. We applied min-max normalization to the pupil-diameter channels in COLET. In addition, both TSFMs apply per-instance normalization before the encoder.

**TABLE I.** Dataset characteristics after downsampling.

| Dataset | Sampling Rate | Total Samples | Class Split % (Low / High) |
| --- | --- | --- | --- |
| EM-COGLOAD | 80 Hz | 8,704 | 49.5 / 50.5 |
| COLET-binary | 120 Hz | 2,061 | 64.1 / 35.9 |

### B. Model Architectures

We studied two TSFMs and two baselines, taking a 512-sample four-channel window as input to each. **MOMENT** (40M params, *d*_*model*_ = 512) is a T5-based, multi-task pre-trained time-series model [14], which we adapted with a custom linear classification head. **Moirai** (14M params, *d*_*model*_ = 384) is a patch-based forecasting model employing any-variate attention [15]; we split the 4 channels into 16 32-length patch segments, projected them into *d*_*model*_, encoded them with the variate and time indices, and pooled the resulting token states by their mean before passing them to a linear classifier head. We benchmarked these against two models that process the window holistically, without sub-patching: a 37K-parameter **CNN baseline** (three 1D Conv-BatchNorm-ReLU-MaxPool layers over the whole window, then average-pooled) and a 567K-parameter **FFN baseline** (2048 *→* 256 *→* 128*→* 64).

### C. Training Procedure

We used subject-level 5-fold cross-validation, partitioning the participants into five folds and excluding all windows from the held-out fold’s subjects during training. For the TSFMs, we applied a two-stage fine-tuning process outlined in Fig. 1. In the first stage, we trained only the linear probe for 15 epochs with the entire backbone frozen; in the second, we tuned targeted backbone layers for up to 25 epochs with early stopping and a patience of 5. We tuned the normalization and positional embeddings for MOMENT, and the final two layers for Moirai. We trained both the CNN and FFN baselines for 40 epochs with early stopping and a patience of 5. All models used a batch size of 256, an Adam optimizer with a learning rate of 0.001, and cross-entropy loss. All scores were evaluated with accuracy, AUC, and macro F1 and 95% CI calculated using a t-distribution of the 5 test set folds.

**Fig. 1.**
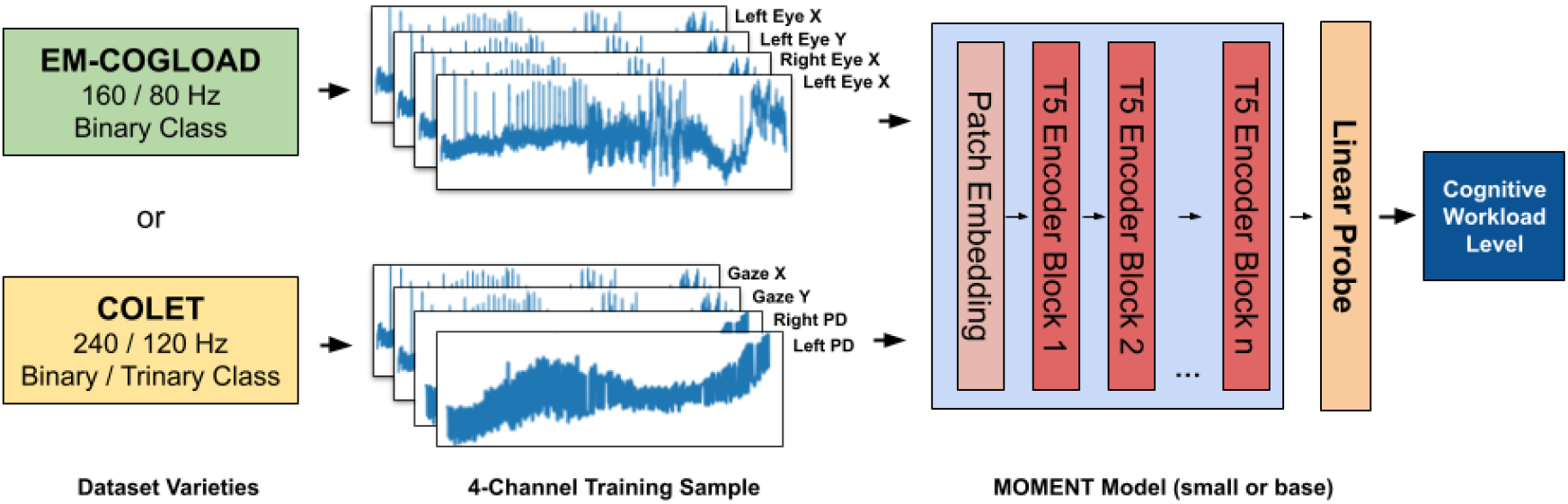
The fine-tuning pipeline for the TSFM models. The selected dataset is converted to four-channel training samples, which are encoded by the pretrained backbone and classified by a linear head. MOMENT uses T5 encoder blocks (6 for small), while Moirai uses a transformer encoder with patch-based input projection.

## III. Results

All models were evaluated under subject-level 5-fold cross-validation, in which all windows from a held-out subject are excluded from training ensuring that every test subject is unseen. This measures generalization to new individuals rather than the memorization of participant-specific ocular characteristics. Table II reports accuracy, AUC, and F1-score averaged across the five folds with 95% confidence intervals, and Fig. 2 shows the per-fold AUC distribution.

**TABLE II.** Subject-level 5-fold cross-validation results. All windows from each test subject are held out during training. *±* values are 95% confidence intervals across folds.

| Model | Params | Acc (%) | AUC | F1 |
| --- | --- | --- | --- | --- |
| *EM-COGLOAD (8,704 windows, 50/50)* |  |  |  |  |
| CNN | 37K | 62.0±9.1 | .693±.109 | .655±.048 |
| FFN | 567K | 67.8±4.4 | .727±.061 | .672±.040 |
| LSTM | 135K | 60.4±5.0 | .698±.048 | .544±.185 |
| MOMENT-sm | 40M | 79.9±3.0 | .882±.029 | .799±.042 |
| **Moirai-sm** | **14M** | **83.9±2.7** | **.918±.023** | **.834±.035** |
| *COLET (2,061 windows, 64/36)* |  |  |  |  |
| CNN | 37K | 62.1±11.5 | .595±.243 | .212±.263 |
| FFN | 567K | 60.2±9.0 | .526±.092 | .369±.211 |
| LSTM | 135K | 63.4±6.6 | .597±.248 | .175±.196 |
| **MOMENT-sm** | **40M** | **63.2±4.3** | **.670±.086** | **.361±.125** |
| Moirai-sm | 14M | 59.9±5.9 | .606±.087 | .416±.102 |

**Fig. 2.**
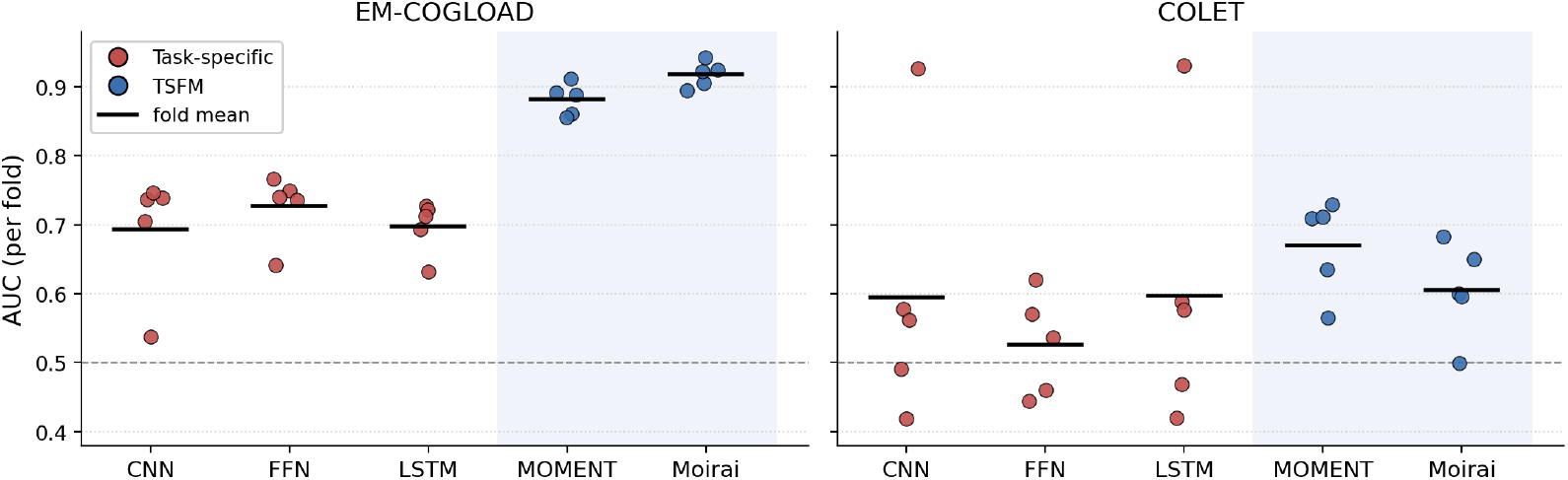
Per-fold subject-level AUC for each model on both datasets. Each point is one of the five cross-validation folds and the black bar is the fold mean. The pretrained TSFMs (shaded) cluster tightly at high AUC on EM-COGLOAD, whereas the task-specific baselines are lower and far more variable across subject folds. On the imbalanced COLET, the CNN and LSTM range from near-chance to above 0.9 depending on which subjects are held out, while MOMENT remains comparatively stable.

### A. EM-COGLOAD Results

On the balanced EM-COGLOAD dataset, the two pre-trained TSFMs generalized markedly better to unseen subjects than the task-specific baselines. Moirai-small achieved the highest performance (83.9% accuracy, 0.918 AUC), followed by MOMENT-small (79.9%, 0.882 AUC), while the CNN, FFN, and LSTM baselines clustered near 0.69–0.73 AUC. The TSFMs were also the most stable across folds, with the narrowest confidence intervals (*±* 0.023 for Moirai and *±* 0.029 for MOMENT), compared to the task-specific models (up to *±*0.109).

### B. COLET Results

On the imbalanced COLET dataset, all models degraded substantially and no model exceeded 0.67 AUC, reflecting the difficulty of the minority (high-workload) class under a 64/36 split. MOMENT was the most robust (63.2% accuracy,0.670 AUC), while the remaining models, including Moirai (0.606 AUC), fell toward chance. The task-specific baselines were also highly unstable, with CNN and LSTM AUC confidence intervals spanning *±* 0.24, indicating their performance depends heavily on which subjects are held out. A decision tree trained on fixation, saccade, blink, and pupil features obtained 74% accuracy and 0.73 F1 [6], indicating feature-engineered approaches remain competitive on this dataset.

### C. Cross-Model Stability

Beyond mean performance, subject-level evaluation reveals differences in stability across models, visible in the per-fold spread in Fig. 2. The pretrained TSFMs are consistently the most stable: on EM-COGLOAD their five-fold results cluster within a narrow band (Moirai spanning only 0.90–0.94 AUC), whereas the task-specific models scatter far more widely. This instability is most pronounced on COLET, where the CNN and LSTM each swing from near-chance on some folds to above 0.9 on others. This indicates that their apparent mean performance depends heavily on which subjects are held out. Pretrained representations remain comparatively uniform and are transferable to unseen users.

## IV. Discussion

In this paper, the two pretrained TSFMs generalize much better to unseen individuals than the task specific baselines. This is most evident on the balanced EM-COGLOAD dataset where Moirai and MOMENT achieve AUCs of 0.918 and 0.882, respectively, while the CNN, FFN, and LSTM are near 0.70. The performance gap suggests that the discriminating features encoded by task specific models are to some degree person dependent, which does not transfer well to new users, whereas TSFMs generalize more uniformly across subjects. Moirai outperforms MOMENT on EM-COGLOAD. This might be attributed to its any-variate attention, which accounts for relationships between channels via explicit bias terms at each feature—a scheme well-suited for multivariate eye-tracking data.

A simple explanation for this gap could be in the nature of the features each model learns. Because task specific models were trained from scratch using data from only several dozen participants, they may learn to predict via shallow, person specific features, such as an individual’s baseline pupil diameter, maximal gaze excursion, or blink frequency, which are useful for a single individual but differ dramatically between people. Such features can appear predictive when the same individuals are seen in training, but they fail to generalize to the held-out subjects in our evaluation, where each person’s own eye signature is unknown. Pretraining TSFMs on varied time-series corpora enables models like those implemented here to acquire abstract temporal dynamics instead, ones which do not depend on any individual person, thereby explaining the higher AUC as well as the dramatically lower variance.

The performance of all models declined on the imbalanced COLET dataset, with only MOMENT performing above chance (AUC = 0.670). Moirai performed comparably to the task-specific baselines, suggesting that the benefits of TSFM pretraining do not generalize uniformly across datasets. In COLET, the combination of severe class imbalance and subjective labels may make the underlying decision boundary less clearly delineated, limiting the ability of pretrained models to learn a robust, generalizable representation. Nevertheless, Moirai exhibited a consistently tighter spread across validation splits (Fig. 2), indicating that when TSFMs do generalize, they may do so more consistently than task-specific models. Because robust generalization is critical for models deployed on users not represented in the training data, these preliminary findings are encouraging. They suggest that TSFMs may have potential for broader deployment in safety-sensitive applications, including driver monitoring, airline crew safety, and medical applications such as assessing surgeon or pilot performance.

Our subject-level evaluation was limited to two datasets with relatively small cohorts of 47 and 52 participants. Larger and more diverse cohorts are needed to validate the generalizability of these initial findings and better characterize model performance across populations and recording conditions. In addition, the task-specific baselines were evaluated using a single run without a fixed random seed; repeating these experiments across multiple seeds would provide more robust estimates of performance variability and confidence. Future work should also investigate parameter-efficient fine-tuning approaches, such as Low-Rank Adaptation (LoRA), to determine whether TSFMs can be efficiently adapted to new tasks while preserving their cross-dataset generalization capabilities. Extending this analysis to models with longer context windows, such as TimesFM, may further improve performance on complex temporal patterns. Finally, reducing TSFM computational cost remains an important priority. Improving inference efficiency could preserve the generalization advantages of these models while reducing latency and enabling deployment in on-device and other time-critical applications.

## V. Conclusion

In this work, we evaluated two time-series foundation models (TSFMs), MOMENT and Moirai, against CNN, FFN, and LSTM baselines for cognitive workload classification from eye-tracking data. Subject-level cross-validation ensured that test subjects remained entirely unseen during training, providing a rigorous assessment of cross-subject generalization. On the class-balanced EM-COGLOAD dataset, both pretrained TSFMs generalized strongly to unseen individuals, with Moirai achieving the best performance (AUC = 0.918) and substantially outperforming the task-specific baselines (*≈* 0.70 AUC). Performance declined on the class-imbalanced COLET dataset, with MOMENT remaining the most robust model (AUC = 0.670). Across both datasets, the TSFMs also demonstrated greater stability across held-out subjects. Collectively, these findings suggest that pretrained TSFMs can provide a substantial advantage over task-specific models when generalizing to unseen individuals in data-limited eye-tracking applications, although the magnitude of this benefit depends on dataset characteristics, particularly class balance and task difficulty. Future work will investigate longer-context architectures such as TimesFM, parameter-efficient fine-tuning methods such as LoRA adapters, and substantially larger and more diverse, class-balanced cognitive workload datasets to further assess the scalability and generalizability of TSFMs.

## Notes

### Competing Interest Statement

The authors have declared no competing interest.

